# ARP2/3 complex mediates the neuropathology of PTEN-deficient human neural cells downstream of mTORC1 and mTORC2 hyperactivation

**DOI:** 10.1101/2025.08.22.670964

**Authors:** Navroop K. Dhaliwal, Octavia Yifang Weng, Ai Tian, Aditi Aggarwal, Mai Ahmed, Guoria Sun, Afrin Bhattacharya, Wendy W.Y. Choi, Haruka Nishimura, Pragnya Chakraborty, Xiaoxue Dong, Yuncheng Wu, Michael D. Wilson, Lu-Yang Wang, Luis F Parada, Julien Muffat, Yun Li

## Abstract

Mutations in the phosphatase and tensin homolog (*PTEN*) gene are linked to severe neurodevelopmental disorders. Loss of *PTEN* causes hyperactivation of both mechanistic target of rapamycin (mTOR) complexes, mTORC1 and mTORC2. Recent studies have shown that this dual hyperactivation is required for the neuropathology observed in *PTEN*-deficient human stem cell-derived neural cells. However, the molecular effectors that integrate these synergistic signals remain unknown. Here, we identify the actin-regulating ARP2/3 complex as a critical point of convergence downstream of mTORC1 and mTORC2. We show that concurrent hyperactivation of both complexes drives increased filamentous actin and elevated levels of the ARP2/3 complex subunits in *PTEN*-deficient human neural precursors and neurons. Pharmacological or genetic inhibition of ARP2/3 is sufficient to rescue multiple disease-relevant phenotypes, including neural precursor hyperproliferation, neuronal hypertrophy, and electrical hyperactivity, without affecting the upstream mTORC1 or mTORC2 hyperactivation. Together, these findings reveal the PTEN-mTOR-ARP2/3 signaling axis as a core mechanism of neuropathology and highlight ARP2/3 inhibition as a potential therapeutic strategy for *PTEN*-related neurodevelopmental disorders.

## Introduction

Germline heterozygous loss-of-function mutations in the *PTEN* gene cause a spectrum of clinical manifestations known as *PTEN* Hamartoma Tumor Syndrome (PHTS), which includes a high prevalence of neurodevelopmental disorders (NDDs) such as autism spectrum disorder, intellectual disability, and epilepsy (1–3). A central mechanism underlying PHTS pathophysiology is the hyperactivation of the phosphoinositide 3-kinase (PI3K)-AKT-mTOR signaling pathway, a master regulator of cell growth, proliferation, and metabolism (4). Consequently, inhibitors of mTOR such as rapamycin have been considered as therapeutic avenues (5). However, clinical trials with mTOR inhibitors have yielded limited efficacy and significant side effects, reflecting the pathway’s broad role in cellular homeostasis (6, 7). These limitations underscore the need to identify more specific downstream effectors that drive neuropathology.

The mTOR kinase serves as the catalytic core of two distinct complexes, mTORC1 and mTORC2 (4, 8). While the roles of these separate complexes have been studied extensively, their interplay in the context of *PTEN*-related NDDs has been less clear. Using human pluripotent stem cells (hPSCs) derived neural models that recapitulate key aspects of the human disorder, we previously demonstrated that synergistic hyperactivation of both mTORC1 and mTORC2 is required to drive the neural phenotypes observed in *PTEN*-deficient human neural precursors (NPs) and neurons, including hyperproliferation, cellular hypertrophy, and electrical hyperactivity (9, 10). Importantly, selective normalization of either mTORC1 or mTORC2, via genetic disruption of *RAPTOR* or *RICTOR*, key components of each respective complex, was sufficient to prevent these abnormalities. These findings suggest that mTORC1 and mTORC2 converge on a common downstream effector or cellular process. Identifying this point of convergence is a critical next step toward developing more targeted therapeutics.

A promising candidate for this convergence is the actin cytoskeleton, a dynamic network of filaments crucial for cell morphology, migration, and synaptic function. Dysregulation of the actin cytoskeleton is recognized as a significant contributor to the pathophysiology of various NDDs (11). For instance, mutations in key actin-related genes, such as *RAC1* and *WASF1* (also known as *WAVE1*) are known to cause NDDs that share clinical manifestations with *PTEN*-related NDDs, including macrocephaly, intellectual disability, epilepsy and autism spectrum disorders (12–14). Furthermore, altered neuronal actin content and dynamics have been mechanistically linked to other syndromic NDDs, including Fragile X syndrome, Tuberous Sclerosis, and Phelan-McDermid Syndrome (15–17). Both mTORC1 and mTORC2 have been independently shown to regulate actin dynamics. mTORC2, for example, is a well-established regulator of small GTPases such as RAC1 and RHOA, which in turn control actin polymerization and cytoskeletal organization (18–20). mTORC1 has also been directly implicated in controlling cytoskeletal organization and filamentous actin (F-actin) content in neuronal contexts (20–22). Together, these insights raise the possibility that actin cytoskeleton may also contribute to the *PTEN*-deficient human neural cells, acting downstream of mTOR hyperactivation.

In this study, we used hPSC-derived neural models to investigate how mTOR hyperactivation drives neuropathology in *PTEN*-deficient cells. We found that dual hyperactivation of mTORC1 and mTORC2 increases levels of the actin-nucleating ARP2/3 complex and elevate F-actin content in both NPs and neurons. Inhibiting ARP2/3, either pharmacologically or genetically, rescued the core cellular and physiological deficits in *PTEN*-mutant human neural cells, including hyperproliferation, neuronal hypertrophy, and electrical hyperactivity. Together, our results establish a direct mechanistic link between mTOR and ARP2/3 signaling axis as the driving mechanism underlying *PTEN*-deficient neuropathology.

## Results

### *PTEN* mutant human neural precursors (NPs) and neurons display increased actin cytoskeleton content driven by mTORC1 and mTORC2 hyperactivation

Actin cytoskeleton dysregulation has been increasingly implicated in neurodevelopmental disorders (NDDs), but its role in *PTEN*-deficient human neuropathology remains poorly understood. To investigate this, we previously used CRISPR/Cas9-mediated gene editing to generate *PTEN* homozygous null mutant hPSCs from WIBR3, a well-characterized female embryonic stem cell line (10). Upon directed differentiation into cortical NPs, these *PTEN* mutant cells lacked PTEN protein and exhibited significantly elevated proliferation compared to isogenic controls, a disease-relevant phenotype that likely contributes to macrocephaly (9, 10). Given that the actin cytoskeleton regulates key cellular processes including cell proliferation and morphology, we first asked whether actin content is altered in *PTEN* mutant NPs. To visualize filamentous actin (F-actin), we performed phalloidin staining. When normalized to soma size, phalloidin intensity was significantly increased in *PTEN* mutant NPs compared to isogenic controls, indicating elevated F-actin content (Figure 1A-1B).

**Figure 1.**
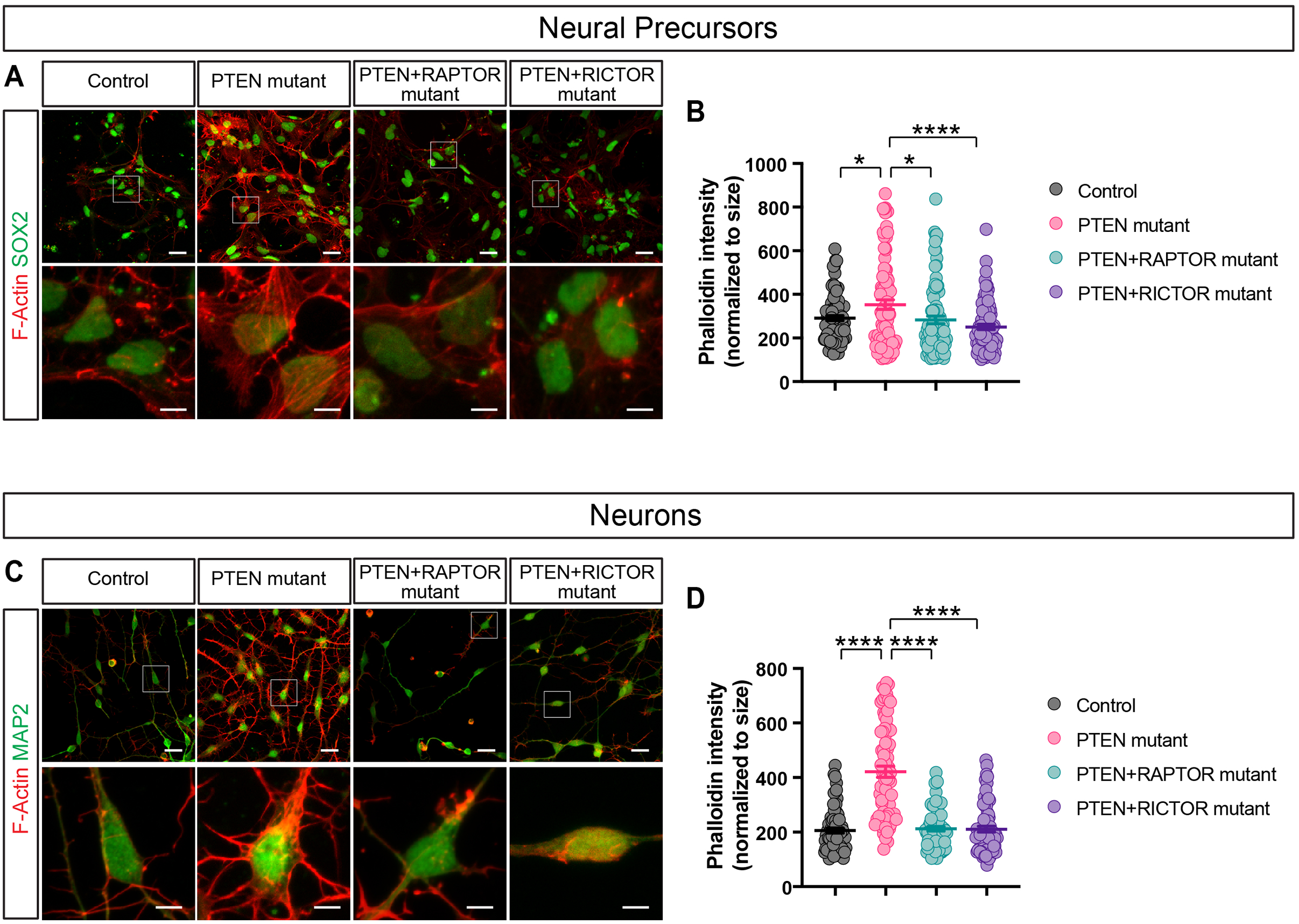
Dual hyperactivation of mTORC1 and mTORC2 drives increased F-actin in *PTEN* mutant human NPs and neurons. (A-B) Representative images (A) and quantification (B) of Phalloidin staining show that *PTEN* mutant WIBR3 human NPs have increased levels of F-actin. Both *PTEN*+*RAPTOR* and *PTEN*+*RICTOR* double mutants, which selectively normalize mTORC1 or mTORC2 signaling, respectively, rescue the F-actin levels. Lower panels are higher magnification views of the boxed areas in the corresponding upper panels. Scale bars in upper panels, 20 μm. Scale bars in lower panels, 5 μm. Each data point represents a cell (n= 82-91 for each group from 4 independent experiments). *p* values are calculated using post hoc multiple comparison after ANOVA. (C-D) Representative images (C) and quantification (D) of Phalloidin staining show that day 30 *PTEN* mutant WIBR3 human neurons have increased levels of F-actin. Both *PTEN*+*RAPTOR* and *PTEN*+*RICTOR* double mutants rescue the F-actin levels. Neurons are induced from WIBR3 NPs using Ngn2 induction. Lower panels are higher magnification views of the boxed areas in the corresponding upper panels. Scale bars in upper panels, 20 μm. Scale bars in lower panels, 5 μm. Each data point represents a cell (n= 70-75 for each group from 4 independent experiments). *p* values are calculated using post hoc multiple comparison after ANOVA. Results are mean +/- SEM. \**p* <0.05, \*\*\*\**p* <0.0001.

We then asked whether this increase in F-actin content observed in *PTEN* mutant NPs is directly caused by mTOR hyperactivation by examination of the role of mTORC1 and mTORC2 individually. Our previous work showed that the synergistic hyperactivation of both mTORC1 and mTORC2 is required for the hyperproliferation phenotypes in *PTEN*-deficient human NPs (9). To test whether this dual activation also underlies actin cytoskeletal dysregulation, we analyzed four genotypes of human NPs: control, *PTEN* mutant, *PTEN+RAPTOR*, and *PTEN+RICTOR* double mutant. In these double mutants, mTORC1 or mTORC2 hyperactivation is selectively rescued by disrupting RAPTOR or RICTOR, key components of mTORC1 and mTORC2, respectively (9). These double mutant NPs were differentiated from clonal hPSC lines carrying *PTEN* homozygous null mutations, along with compound heterozygous mutations in *RAPTOR* or *RICTOR* (consisting of one frameshift and one frame-preserving mutation) that reduce their protein levels by approximately 50% (9). Phalloidin staining showed that disruption of either *RAPTOR* or *RICTOR* restored F-actin levels to those observed in controls (Figure 1A-1B). These findings demonstrate that the simultaneous hyperactivation of both mTORC1 and mTORC2 is required for elevated actin content in *PTEN* mutant human NPs.

We next investigated whether similar cytoskeletal abnormalities occur in *PTEN* mutant human cortical neurons. Using a doxycycline-inducible Ngn2 overexpression lentivirus, we transduced control, *PTEN*, *PTEN+RAPTOR*, and *PTEN+RICTOR* mutant WIBR3 hPSCs and differentiated them into neurons. Phalloidin staining revealed significantly increased F-actin content, even after normalizing to cell size (Figure 1C-1D). This phenotype was rescued in both *PTEN+RAPTOR* and *PTEN+RICTOR* double mutant neurons, indicating that dual mTORC1 and mTORC2 hyperactivation is required for actin cytoskeleton dysregulation in *PTEN*-deficient neurons (Figure 1C-1D), as in NPs.

To validate these findings in an independent genetic background, we used CRISPR/Cas9 to generate *PTEN* homozygous null mutant hPSC lines in the PGPC17-iNgn2 system (Figure S1B-S1C). PGPC17 is a well-characterized hPSC line reprogrammed from a healthy male donor (23), and the PGPC17-iNgn2 version contains a doxycycline-inducible Ngn2 expression cassette in the *CLYBL* safe-harbor locus (Figure S1A). Upon doxycycline treatment, isogenic control and *PTEN* mutant hPSCs were differentiated into cortical neurons. Phalloidin staining confirmed that *PTEN* mutant neurons in this independent genetic background also exhibit significantly increased F-actin content (Figure S2A-S2B).

### Synergistic mTORC1 and mTORC2 hyperactivation increase levels of the ARP2/3 complex in *PTEN* mutant human NPs and neurons

To investigate the molecular mechanisms of increased F-actin content in *PTEN* mutant WIBR3 human neural cells, we examined the protein levels of key regulators of actin dynamics. In both *PTEN* mutant NPs and neurons, we observed elevated levels of RAC1, a small GTPase known to be a key organizer of the actin cytoskeleton (Figure 2A-2B, 2G-2H). This finding is consistent with previous studies in *Pten* null mutant mouse fibroblasts (24). Given the broad role of RAC1 in regulating cytoskeleton dynamics, we sought to further specify this mechanism. RAC1 is known to regulate the ARP2/3 complex, a critical actin nucleator that generates branched actin networks and plays important roles in neurodevelopment (25, 26). We found that *PTEN* mutant NPs and neurons had significantly elevated levels of multiple ARP2/3 subunits, including ARP2, ARP3, ARPC1, and ARPC4 (Figure 2A-2L).

**Figure 2.**
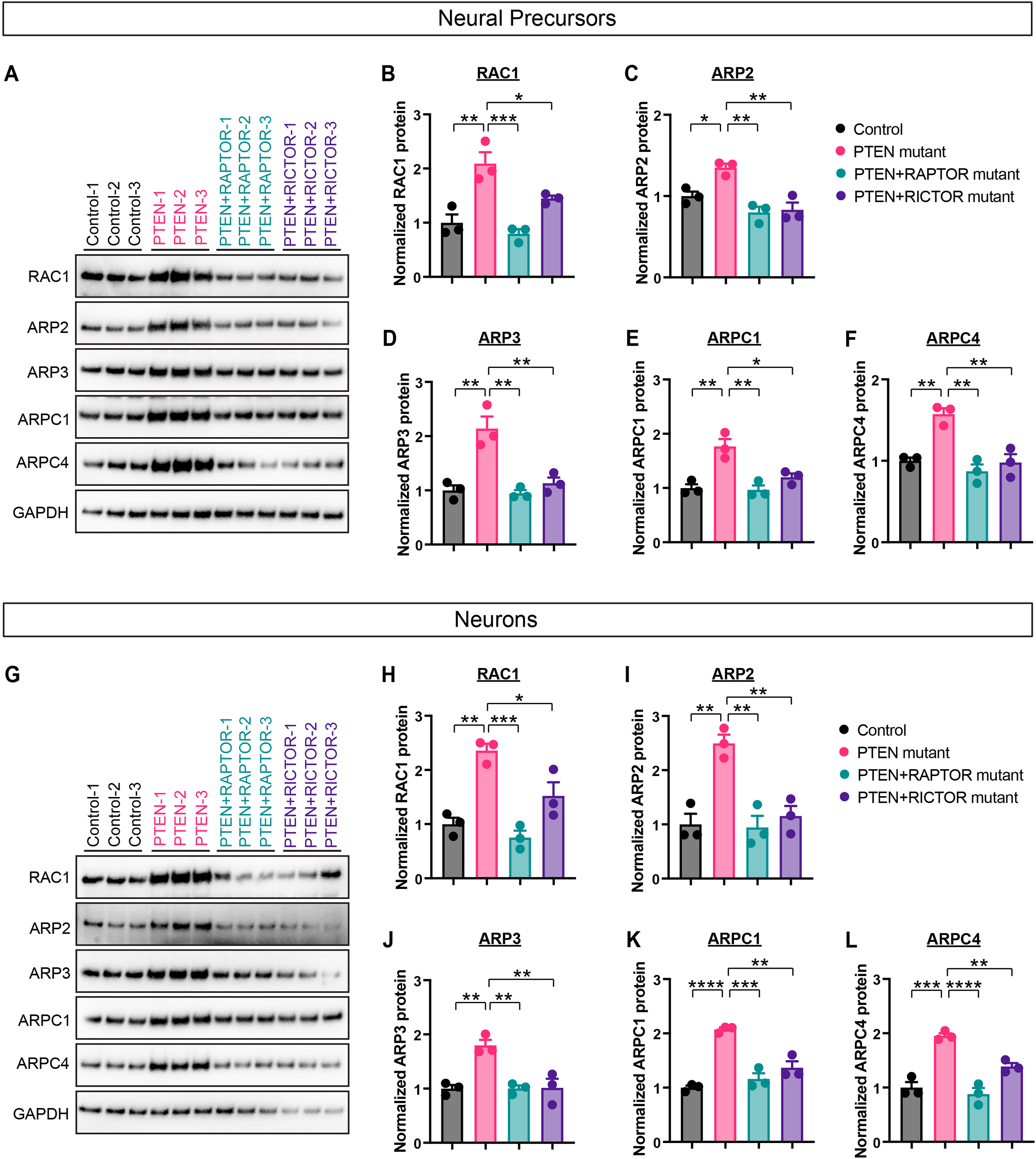
Dual hyperactivation of mTORC1 and mTORC2 causes increased levels of the ARP2/3 complex in *PTEN* mutant human NPs and neurons. (A-F) Immunoblotting analysis shows that *PTEN* mutant WIBR3 human NPs have increased levels of RAC1 and key components of the ARP2/3 complex (ARP2, ARP3, ARPC1, and ARPC4). Both *PTEN*+*RAPTOR* and *PTEN*+*RICTOR* double mutants rescue these protein levels. Each data point represents an independent experiment (different hPSC clone or independent differentiation, n=3 for each group). *p* values are calculated using post hoc multiple comparison after ANOVA. (G-H) Immunoblotting analysis shows that day 10 *PTEN* mutant WIBR3 human neurons have increased levels of RAC1 and key components of the ARP2/3 complex (ARP2, ARP3, ARPC1, and ARPC4), which are rescued in both *PTEN*+*RAPTOR* and *PTEN*+*RICTOR* double mutants. Neurons are induced from WIBR3 NPs using Ngn2 induction. Each data point represents an independent experiment (different hPSC clone or independent differentiation, n=3 for each group). *p* values are calculated using post hoc multiple comparison after ANOVA. Results are mean +/-SEM. \**p* <0.05, \*\**p* <0.01, \*\*\**p* <0.005, \*\*\*\**p* <0.0001.

To determine whether RAC1 and ARP2/3 upregulation depends on mTORC1 and mTORC2 hyperactivation, we analyzed protein levels in *PTEN+RAPTOR* and *PTEN+RICTOR* double mutant human NPs and neurons. In both genotypes, the elevated levels of ARP2/3 subunits and RAC1 were rescued to control levels, indicating that both pathways are required, and further supporting the model in which the mTORC1 and mTORC2 hyperactivation synergistically regulate actin cytoskeleton (Figure 2A-2L). Lastly, we confirmed that ARP2/3 complex components are also significantly elevated in *PTEN* homozygous null mutant neurons derived from the PGPC17 hPSC line (Figure S2C-S2D).

### Pharmacological inhibition of the ARP2/3 complex rescues *PTEN* mutant NPs and neurons

We next asked whether the ARP2/3 complex is required for the disease-relevant molecular, cellular, and physiological phenotypes of *PTEN* mutant human NPs and neurons. We treated control and *PTEN* mutant NPs with the ARP2/3 inhibitor CK-666 (27, 28) (100μM) for 24 hours and observed significantly reduced F-actin content as measured by phalloidin staining (Figure 3A-3B). Using KI67 immunostaining and an EdU incorporation assay, we showed that *PTEN* mutant human NPs display hyperproliferation, consistent with our previous reports (9, 10). The ARP2/3 inhibitor CK-666 was effective in reducing the number of proliferating cells to control level (Figure 3C-3D).

**Figure 3.**
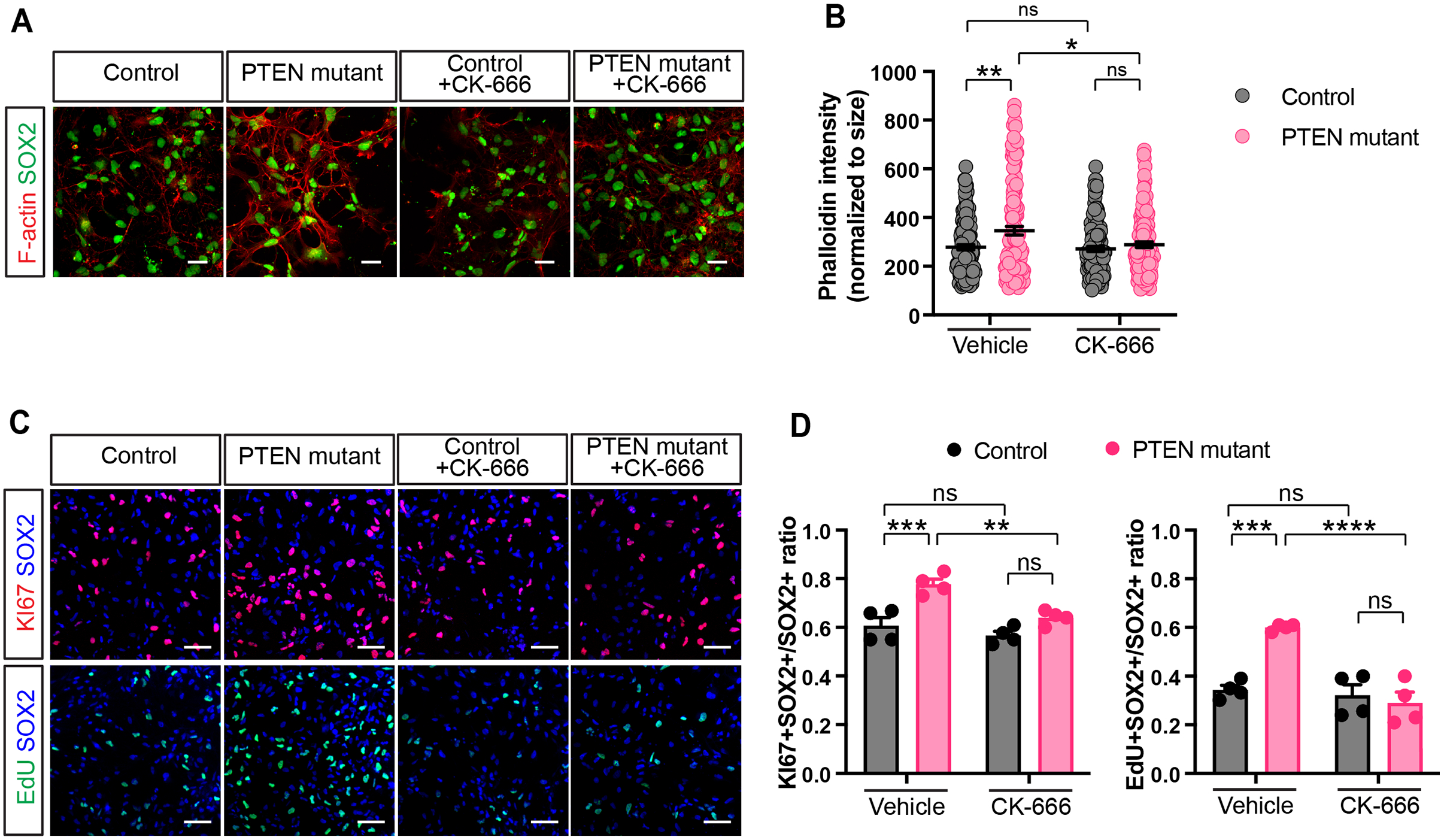
The ARP2/3 complex inhibitor CK-666 restores F-actin levels and rescues hyperproliferation in *PTEN* mutant human NPs. (A-B) Representative images (A) and quantification (B) show that chronic CK-666 treatment reduces the increased levels of F-actin in *PTEN* mutant WIBR3 human NPs. Scale bars, 20 μm. Each data point represents a cell (n= 115-131 for each group from 4 independent experiments). *p* values are calculated using post hoc multiple comparison after ANOVA. (C-D) Representative images (C) and quantification (D) show that chronic CK-666 treatment reduces the over-proliferation of *PTEN* mutant WIBR3 human NPs, as measured by Ki67 immuno-staining and EdU incorporation assay. Scale bar, 50 μm. Each data point represents an independent experiment (different hPSC clone or independent differentiation, n= 4 for each group). *p* values are calculated using post hoc multiple comparison after ANOVA. Results are mean +/-SEM. \**p* <0.05, \*\**p* <0.01, \*\*\**p* <0.005, \*\*\*\**p* <0.0001.

Next, we treated differentiating control and *PTEN* mutant PGPC17 neurons with the CK-666 inhibitor (100μM) starting on day 1 of Ngn2 induction and continuously throughout. Phalloidin staining showed that chronic CK-666 treatment significantly reduced the F-actin content in *PTEN* mutant neurons on day 14 (Figure 4A-4B). While *PTEN* mutant neurons showed significant increases in soma and nucleus sizes as well as dendritic arborization, CK-666 treatment rescued these phenotypes to control level (Figure 4C-4G). We recorded neuronal electrical activity using a multi-electrode array (MEA) platform over the span of 30 days. *PTEN* mutant neurons were significantly more active and synchronized in their firing pattern compared to control neurons, consistent with our previous report (9). Chronic CK-666 treatment prevented the development of these hyperactive electrical phenotypes (Figure 4H-4J, Video S1). Together, these findings demonstrate that the ARP2/3 complex mediates the disease-relevant morphological and physiological phenotypes in *PTEN* mutant human NPs and neurons.

**Figure 4.**
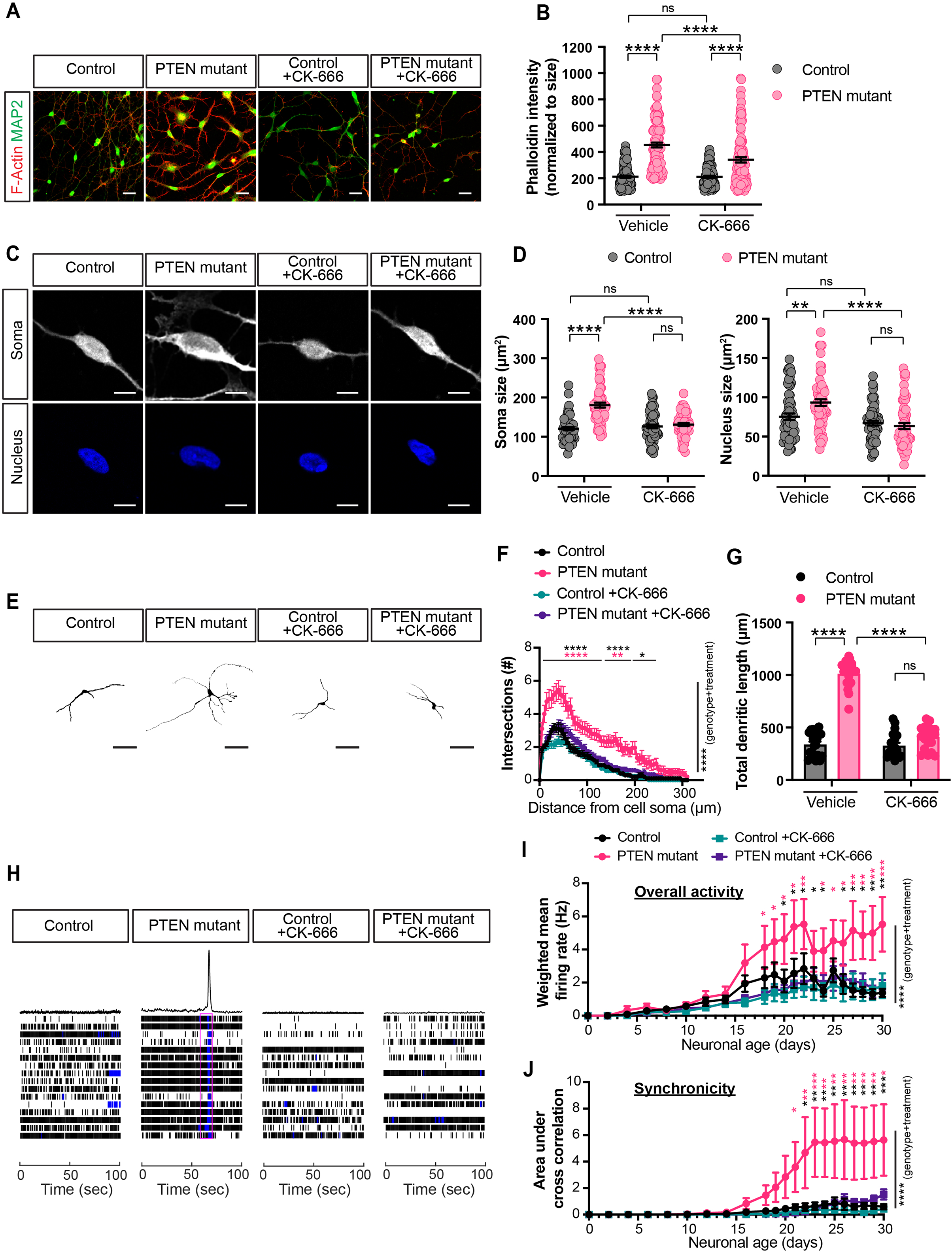
The ARP2/3 complex inhibitor CK-666 restores F-actin level and rescues morphological and electrophysiological phenotypes in *PTEN* mutant human neurons. (A-B) Representative images (A) and quantification (B) show that chronic CK-666 treatment reduces the increased levels of F-actin in *PTEN* mutant PGPC17 human neurons on day 14. Scale bars, 20 μm. Each data point represents a cell (n= 101-105 for each group from 3 independent experiments). *p* values are calculated using post hoc multiple comparison after ANOVA. (C-D) Representative images (C) and quantification (D) show that chronic CK-666 treatment reduces the soma and nucleus size of *PTEN* mutant human neurons on day 14. Scale bar, 10 μm. Each data point represents a cell (n= 57-61 for each group from 3 independent experiments). *p* values are calculated using post hoc multiple comparison after ANOVA. (E) Representative images of the traces of individual GFP-labeled neurons showing their dendritic morphology on day 14. Scale bar, 50 μm. (F-G) Sholl analyses reveal increased dendritic complexity (F) and total dendritic length (G) in *PTEN* mutant human neurons, which are rescued by chronic CK-666 treatment. ANOVA found significant effects of genotype and treatment (F, black asterisks marked on the right side of the graph). Post hoc multiple comparison test revealed differences at specific distance from cell soma, in which the black asterisk represents comparison between control and *PTEN* mutant neurons, and the red asterisk represents comparison between *PTEN* mutant neurons treated with vehicle or CK-666 (F). Each data point represents a cell (n=20 for each group from 3 independent experiments). *p* values are calculated using post hoc multiple comparison after ANOVA. (H-J). Multi-electrode array analysis shows that chronic CK-666 treatment rescues the electrical hyperactivity (I, mean firing rate) and hyper-synchronization (J) phenotypes of *PTEN* mutant human neurons. Raster plots (H) show spike distribution on day 30. The purple rectangle highlights the incidence of network burst. ANOVA found significant effects of genotype and CK-666 treatment (black asterisks marked on the right side of the graphs). Post hoc multiple comparison test revealed differences on individual days of recording, in which the black asterisk represents a comparison between vehicle-treated control and *PTEN* mutant neurons, and the red asterisk represents a comparison between vehicle- and CK-666-treated *PTEN* mutant neurons. N= 6 for each group from 6 independent experiments. Results are mean +/-SEM. \**p* <0.05, \*\**p* <0.01, \*\*\**p* <0.005, \*\*\*\**p* <0.0001.

### Genetic inhibition of the ARP2/3 complex rescues *PTEN* mutant NPs and neurons

We sought to further validate the role of the ARP2/3 complex using a CRISPR-based genetic approach to disrupt *ARPC4*, an essential subunit of the ARP2/3 complex. Interestingly, patients with *ARPC4* germline loss-of-function mutations develop a rare NDD called DEVLO (Developmental Delay, Language Impairment, and Ocular Abnormalities) (29), which is often characterized by microcephaly, a phenotype opposite of the macrocephaly found in *PTEN*-related NDDs (2), and consistent with the microcephalic phenotypes of a patient with *PTEN*-duplication (30). We generated *PTEN+ARPC4* double mutant WIBR3 NPs in a pooled manner by transducing *PTEN* homozygous null mutant WIBR3 NPs with lentiviruses expressing Cas9 and sgRNA against the exon 2 of *ARPC4*. This polyclonal population of transduced cells were selected with puromycin and assessed via Sanger sequencing to confirm partial *ARPC4* genetic disruption (Figure S3). We further demonstrated that ARPC4 protein levels were significantly reduced compared to *PTEN* single mutants, returning to levels comparable to wild-type controls (Figure 5A-5B). We also observed that other ARP2/3 complex subunits were significantly reduced to control levels, consistent with the knowledge that reduction of ARPC4 disrupts ARP2/3 complex formation (Figure 5C). Phalloidin staining demonstrated that *PTEN+ARPC4* mutant human NPs displayed rescued levels of F-actin content (Figure 5D-5E). We performed KI67 immunostaining and EdU incorporation assay to show that *PTEN+ARPC4* mutant human NPs displayed rescued proliferation levels comparable to isogenic control NPs (Figure 5F-H).

**Figure 5.**
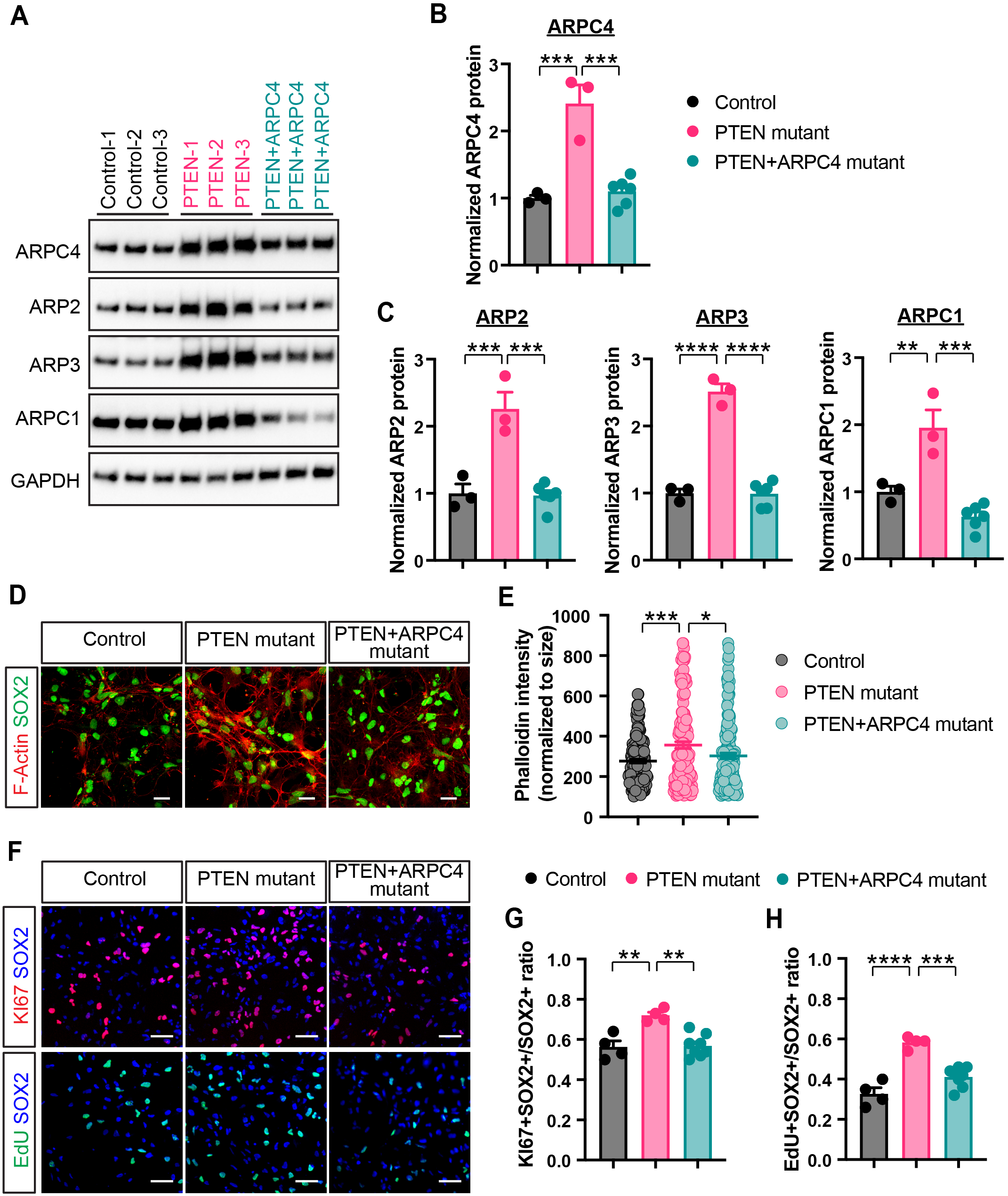
Genetic inhibition of the ARP2/3 complex rescues *PTEN* mutant human NPs. (A-C) Immunoblotting analysis shows reduced ARPC4 protein levels in the *PTEN+ARPC4* double-mutant WIBR3 human NPs. Note the concomitant reduction in other key components of the ARP2/3 complex (ARP2, ARP3, and ARPC1) in the *PTEN+ARPC4* double-mutant WIBR3 human NPs. Each data point represents an independent experiment (different hPSC clone, independent transduction, or ARPC4-sgRNA, n=3-6 for each group). *p* values are calculated using post hoc multiple comparison after ANOVA. (D-E) Representative images (D) and quantification (E) show that *PTEN+ARPC4* double-mutants rescue F-actin level in human NPs. Scale bars, 20 μm. Each data point represents a cell (n= 153-168 for each group from 3-6 independent experiments). *p* values are calculated using post hoc multiple comparison after ANOVA. (F-H) Representative images (F) and quantification (G-H) show that *PTEN+ARPC4* double-mutants rescue the overproliferation phenotypes of *PTEN* mutant human NPs, as measured by Ki67 immunostaining and EdU incorporation assay. Scale bars, 50 μm. Each data point represents an independent experiment (different hPSC clone, independent transduction, or *ARPC4*-sgRNA, n=4-8 for each group). *p* values are calculated using post hoc multiple comparison after ANOVA. Results are mean +/-SEM. \**p* <0.05, \*\**p* <0.01, \*\*\**p* <0.005, \*\*\*\**p* <0.0001.

Next, we performed similar genetic inhibition experiment on human neurons. We transduced *PTEN* homozygous null mutant PGPC17 hPSC lines with sgRNA targeting *ARPC4* to generate a pooled population of *PTEN+ARPC4* mutant hPSCs. Sanger sequencing and immunoblotting confirmed *ARPC4* genetic disruption (Figure S4) and protein reduction in ARPC4 as well as other ARP2/3 complex subunits (Figure 6A-6C). Phalloidin staining showed that *PTEN+ARPC4* mutant neurons displayed rescued F-actin content (Figure 6D-6E), soma and neurite morphology (Figure 6F-6K) and electrical activity (Figure 6L-6N, Video S2).

**Figure 6.**
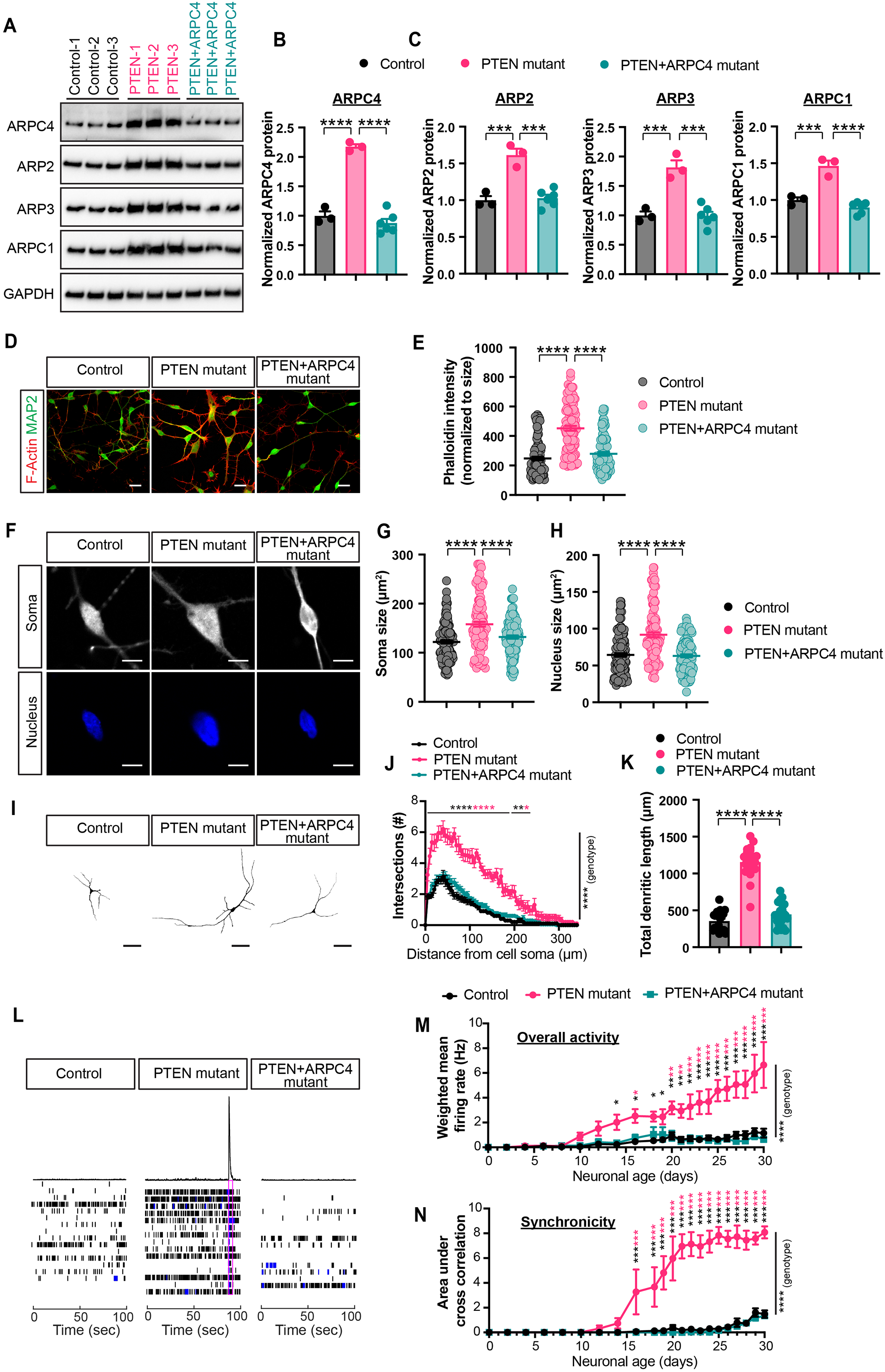
Genetic inhibition of the ARP2/3 complex rescues *PTEN* mutant human neurons. (A-C) Immunoblotting analysis shows reduced ARPC4, as well as other key components of the ARP2/3 complex (ARP2, ARP3, and ARPC1) in the *PTEN+ARPC4* double-mutant PGPC17 human neurons on day 10. Neurons are induced from PGPC17 hPSCs using Ngn2 induction. Each data point represents an independent experiment (different hPSC clone, independent transduction, or *ARPC4*-sgRNA, n=3-6 for each group). *p* values are calculated using post hoc multiple comparison after ANOVA. (D-E) Representative images (D) and quantification (E) show that *PTEN+ARPC4* double-mutants rescues F-actin levels in day 30 human neurons. Scale bars, 20 μm. Each data point represents a cell (n=101-106 for each group from 3-6 independent experiments). *p* values are calculated using post hoc multiple comparison after ANOVA. (F-H) Representative images (C) and quantification (D) show that *PTEN+ARPC4* double-mutants rescues the soma and nucleus size of *PTEN* mutant human neurons on day 30. Scale bar, 10 μm. Each data point represents a cell (n=127-130 for each group from 3-6 independent experiments). *p* values are calculated using post hoc multiple comparison after ANOVA. (I-K) Sholl analysis shows that *PTEN+ARPC4* double-mutants rescue the increased dendritic complexity and total dendritic length in *PTEN* mutant human neurons on day 30. ANOVA found significant effects of genotype and treatment (J, black asterisks marked on the right side of the graph). Post hoc multiple comparison test revealed differences at specific distance from cell soma, in which the black asterisk represents comparison between control and *PTEN* mutant neurons, and the red asterisk represents comparison between *PTEN* and *PTEN+ARPC4* mutant neurons (K). Each data point represents a cell (n=20 for each group from 3-6 independent experiments). *p* values are calculated using post hoc multiple comparison after ANOVA. (L-N). Multi-electrode array analysis shows that *PTEN+ARPC4* double-mutants rescues the electrical hyperactivity (M) and hyper-synchronization (N) phenotypes of *PTEN* mutant human neurons. Raster plots (L) show spike distribution on day 30. The purple rectangle highlights the incidence of network burst. ANOVA found significant effects of genotype (black asterisks marked on the right side of the graphs). Post hoc multiple comparison test revealed differences on individual days of recording, in which the black asterisk represents a comparison between control and *PTEN* mutant neurons, and the red asterisk represents a comparison between *PTEN* and *PTEN+ARPC4* mutant neurons. N= 4 for each group from 4 independent experiments. Results are mean +/-SEM. \**p* <0.05, \*\**p* <0.01, \*\*\**p* <0.005, \*\*\*\**p* <0.0001.

Together, these results corroborate our findings using ARP2/3 inhibitor and demonstrate that ARP2/3 is necessary for the development of cytoskeletal and functional abnormalities in *PTEN* mutant human neural precursors and neurons.

### ARP2/3 inhibition rescues *PTEN* mutant neural phenotypes without normalizing mTORC1 and mTORC2 hyperactivation

We showed that the increased ARP2/3 levels required the concomitant hyperactivation of mTORC1 and mTORC2. To place the ARP2/3 complex within this signaling cascade, we investigated whether ARP2/3 inhibition affects the upstream hyperactivation of mTORC1 and mTORC2. Immunoblotting showed that CK-666 treatment did not rescue the hyperactivation of mTORC1 and mTORC2 in *PTEN* mutant NPs (Figure 7A-7C) and neurons (Figure 7G-7I). Similarly, hyperactivation in these pathways persisted in *PTEN+ARPC4* double mutant NPs and neurons (Figure 7D-F, 7J-L). Taken together, these findings support the model that ARP2/3 is a convergent downstream effector of mTORC1 and mTORC2 pathways (Figure 7M).

**Figure 7.**
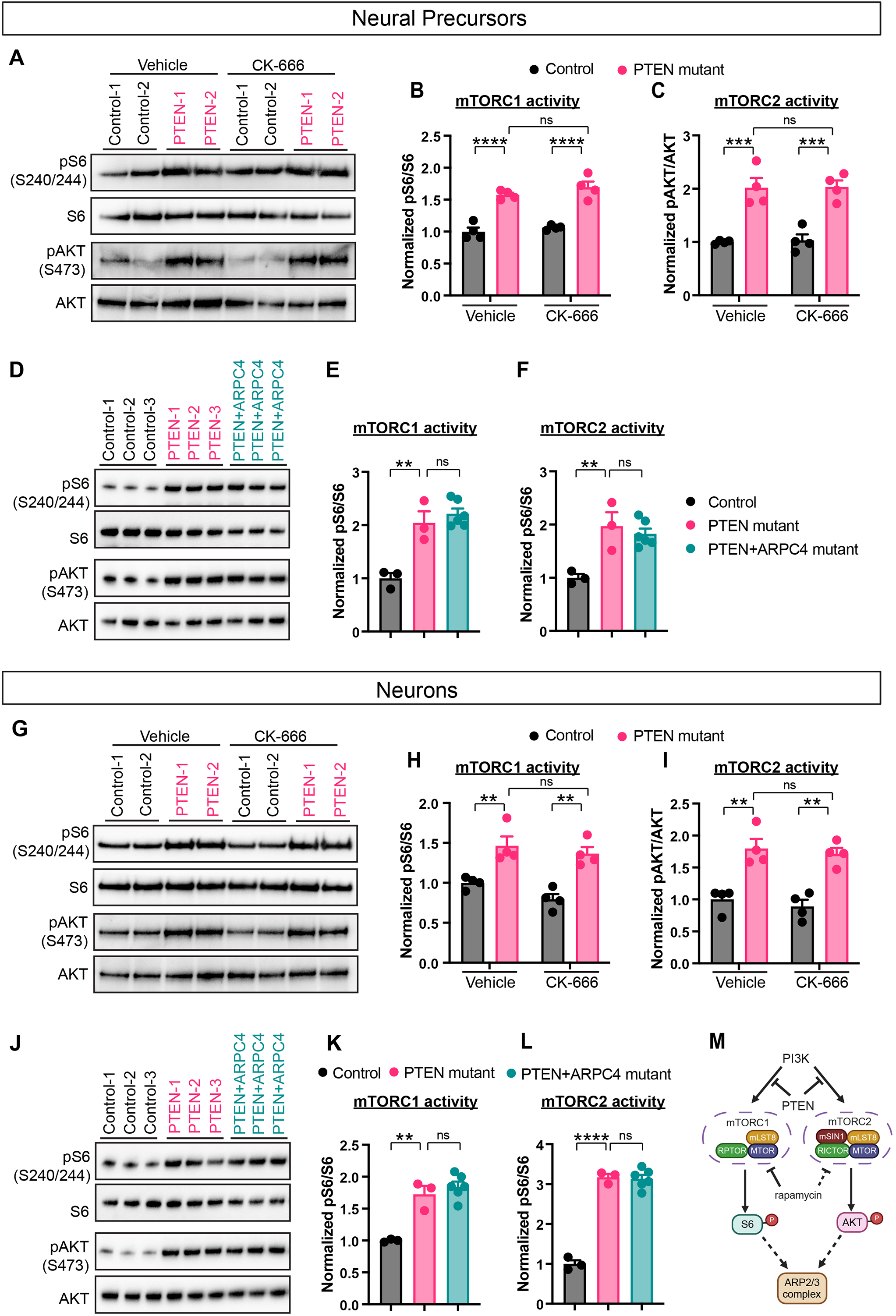
ARP2/3 complex inhibitions rescue *PTEN* mutant neural phenotypes without normalizing mTORC1 and mTORC2 hyperactivation. (A-F) Immunoblotting analysis shows chronic CK666 treatment (A-C) and *PTEN+ARPC4* double-mutants (D-F) did not reduce the hyperactivation of mTORC1 (pS6) and mTORC2 (pAKT) in *PTEN* mutant WIBR3 human NPs. Each data point represents an independent experiment (different hPSC clone, independent differentiation, or *ARPC4*-sgRNA, n=4 for each group of CK-666 inhibition, n=3-6 for each group of genetic inhibition). *p* values are calculated using post hoc multiple comparison after ANOVA. (G-L) Immunoblotting analysis shows chronic CK666 treatment (G-I) and *PTEN+ARPC4* double-mutants (J-L) did not reduce the hyperactivation of mTORC1 (pS6) and mTORC2 (pAKT) in *PTEN* mutant PGPC17 human neurons on day 10. Neurons are induced from PGPC17 hPSCs using Ngn2 induction. Each data point represents an independent experiment (different hPSC clone, independent differentiation, or *ARPC4*-sgRNA, n=4 for each group of CK-666 inhibition, n=3-6 for each group of genetic inhibition). *p* values are calculated using post hoc multiple comparison after ANOVA. (M) Schematic diagram of the PTEN-mTOR-ARP2/3 signaling pathway. Results are mean +/-SEM. \*\**p* <0.01, \*\*\**p* <0.005, \*\*\*\**p* <0.0001.

## Discussion

*PTEN*-deficiency disorder belongs to a wide spectrum of NDDs known as mTORopathies, which are characterized by mTOR hyperactivation and overlapping clinical features, including macrocephaly, epilepsy, and autism (31, 32). Although mTOR hyperactivation is a hallmark of *PTEN*-associated NDDs, the downstream mechanisms linking this signaling abnormality to neural dysfunction remain poorly defined. Our previous work demonstrated that synergistic activation of both mTORC1 and mTORC2 is required to drive the morphological and electrophysiological phenotypes of *PTEN*-deficient human neural cells (9). In the current study, we identify the ARP2/3 complex as a downstream effector of mTORC1 and mTORC2 hyperactivation. Using isogenic hPSC models, we show that *PTEN* loss leads to a marked increase in F-actin content in both NPs and neurons, accompanied by elevated protein levels of RAC1 and multiple ARP2/3 subunits. Both pharmacological inhibition and genetic disruption of ARP2/3 rescued F-actin accumulation, precursor hyperproliferation and neuronal hypertrophy, and prevented the emergence of electrical hyperactivity in neurons. Hyperactivation of both mTORC1 and mTORC2 is required to drive these phenotypes, and ARP2/3 inhibition rescued them without affecting the elevated mTORC1 or mTORC2 activity, positioning ARP2/3 as a downstream effector of this dual pathway activation.

Our findings contribute to a growing recognition that cytoskeletal dysregulation is a key mechanistic driver of NDDs (11, 33). Previous studies in human patients have shown that loss-of-function mutations in ARP2/3 complex components, such as *ARPC4*, cause severe neurodevelopmental abnormalities, including microcephaly and developmental delays (29). Similarly, genetic ablation of ARP2/3 components in conditional knockout mouse models leads to profound defects in neuronal migration, dendritic branching, and synaptic development (34–36). These loss-of-function phenotypes emphasize the essential roles of ARP2/3 during key early stages of neurodevelopment. In contrast to this established literature, our work highlights the detrimental impact of increased ARP2/3 activity in human neural cells. To our knowledge, this is the first demonstration that elevated ARP2/3 levels can contribute to neurodevelopmental pathology. This finding positions the ARP2/3 complex as a dosage-sensitive causal mechanism for NDDs, similar to other genes such as *PTEN, MECP2*, *SHANK3*, *SCN2A*, *UBE3A*, and *RAC1*. Supporting this concept, activating mutations in *RAC1*, a key regulator of ARP2/3, cause macrocephaly in human patients, likely by driving excessive ARP2/3 activation (12). Furthermore, increased gene dosage of *CYFIP1*, a regulator of RAC1, is thought to contribute to the macrocephaly, autism and epilepsy phenotypes in 15q11-13 duplication syndromes (37). It is possible that elevated ARP2/3 activity enhances aberrant actin remodeling and neurite growth, leading to the electrical hyperactivity and increased network synchrony observed in *PTEN* mutant cultures. Future studies to systematically investigate the pathological impact of excessive ARP2/3 complex levels, alone or in the context of other NDDs, could yield further mechanistic insights and therapeutic opportunities.

The current study establishes a causal link from mTOR hyperactivation to elevated F-actin content and ARP2/3 levels. We showed that genetic inhibition of either mTORC1 or mTORC2, via the disruption of *RAPTOR* or *RICTOR*, respectively, is sufficient to rescue the F-actin content and ARP2/3 subunit levels. Although this specific signaling axis has not been previously demonstrated in the context of neurodevelopment, evidence from other biological systems supports our conclusion. For example, it has been reported that F-actin content is pathologically elevated in the aging drosophila brain, which can be rescued by the mTOR inhibitor rapamycin (22). Furthermore, in a mouse model of KRAS-driven pancreatic cancer that displays mTOR hyperactivation, elevated ARP2/3 level was observed and proven relevant as a therapeutic target to reduce tumor progression (28). Using an *in vitro* model of *PTEN*-deficiency disorder, our current study reveals that the mTOR-ARP2/3 axis is a pathogenic driver during human neurodevelopment. Given that mutations in other mTORopathy genes such as *PIK3CA*, *AKT1/3*, and *MTOR* also can activate both mTORC1 and mTORC2, it is possible that ARP2/3 dysfunction represents a shared downstream mechanism contributing to neuropathology across multiple mTOR-driven NDDs.

Our study demonstrates that inhibiting ARP2/3 can ameliorate disease-relevant phenotypes in an *in vitro* model of *PTEN*-deficiency NDD. This aligns with previous work in the aging drosophila brain, where reducing elevated F-actin level via siRNA slowed aging and extended lifespan (22). Although ARP2/3 inhibitors such as CK-666 (used in the current study) have been extensively tested and shown to suppress tumor cell migration and invasiveness in preclinical cancer models (28, 38–40), clinical translation of these inhibitors has yet to occur. Given the importance of ARP2/3 in fundamental cellular processes, including neuronal morphogenesis and synaptic function, direct therapeutic targeting of this complex requires careful consideration. Moreover, it is likely that ARP2/3 contributes to a subset, but not the entirety, of the neural phenotypes associated with PTEN deficiency. Therefore, its inhibition may offer only partial correction of the broad pathological spectrum. A cautious yet plausible direction for future work could involve evaluating combination therapies of inhibitors against mTOR, ARP2/3 or other related downstream effectors. Taken together, our findings support the broader therapeutic concept that targeting downstream effectors of mTOR signaling, such as ARP2/3, may offer a more specific and effective strategy for *PTEN*-related and other mTOR-driven NDDs.

## Acknowledgements

The authors thank YoungJun Ju, Fatima Naimi for technical support, and all members of the Li and Muffat labs for helpful discussions. CLYBL-TO-hNGN2-BSD-mApple was a gift from Michael Ward (Addgene #124229). The PGPC17 cell line was generously provided by James Ellis. This work was supported by grants from the CIHR (PJT-180565), Can-GARD, Epilepsy Canada, and the Scottish Rite Charitable Foundation of Canada. J.M. received funding from the Canada Research Chairs program.

## Author Contribution

N.K.D., O.Y.W., J.M. and Y.L. conceived the project, designed the experiments, interpreted the results, and wrote the paper with input from all authors. N.K.D. and O.Y.W. conducted the experiments and performed data analyses. A.T. generated key reagent. A.A., M.A., G.S., A.B., W.W.Y.C., H.N., X.D., P.C. assisted with experiments. Y.C., M.D.W., L.Y.W. and L.F.P. provided key reagents, supervision, and advised on the study design.

## Materials and Method

### Human embryonic stem cell culture

Human male induced pluripotent stem cell line PGPC17 and female embryonic stem cell line WIBR3 were reported in previous studies (23, 41). Control, *PTEN* mutant, *PTEN+RAPTOR* double mutant, and *PTEN+RICTOR* double mutant WIBR3 hPSCs were generated previous studies (9, 10). hPSCs were cultured on Matrigel (Corning #354234)-coated culture dishes in mTeSR plus medium (Stem Cell Technologies #05825) or iPS brew (Miltenyi Biotec #130-104-368). Cultures were passaged every 5-7 days either manually, or with ReLeSR (Stem Cell Technologies #05873). hPSCs were cryo-preserved in freezing media containing 90% Knock-out Serum Replacement (Thermo Fisher Scientific #10828-028), 10% DMSO (Sigma #D2650), and 10μM Rock Inhibitor Y27632 (Millipore Sigma #146986-50-7). All cell lines were routinely tested for mycoplasma negativity.

### Statement of compliance with IRBs

The PGPC17 and WIBR3 hPSC lines were approved for use by the Stem Cell Oversight Committee of the Canadian Institutes of Health Research, and the Research Ethics Board of the Hospital for Sick Children.

#### Lentivirus production

Lentivirus plasmids containing sgRNA against *ARPC4* were cloned into the PLCKO plasmid (Addgene #73311) according to the published protocol (42). Two different sgRNAs were used (Table S2). The pLV-TetO-hNGN2-Neo inducible Ngn2 lentivirus plasmid was previously reported (43) (Addgene #99378). To generate VSVG-coated lentivirus particles, HEK293 cells were transfected using X-tremeGENE 9 (Sigma #6365809001), with a mixture of lentiviral construct and third-generation packaging plasmids. The culture medium was changed 12 hours after transfection and collected 96 hours after transfection. Virus-containing medium was filtered through a 0.45μm filter and concentrated via ultracentrifugation using Beckman Coulter ultracentrifuge at 23,000rpm and 4°C.

#### CRISPR/Cas9-mediated gene editing in hPSC line

PGPC17-iNgn2 hPSCs were generated via CRISPR/Cas9-mediated genome editing, using sgRNA against the *CLYBL* genomic locus and the CLYBL-TO-hNGN2-BSD-mApple donor plasmid (Addgene #124229), following previously published protocol (44). To generate clonal *PTEN* homozygous mutant PGPC17-iNgn2 hPSCs, CRISPR/Cas9-mediated genome editing was performed as previously described with modifications (10). Briefly, PGPC17-iNgn2 hPSCs were cultured in 10μM Y27632 for 24 hr, harvested using accutase, and electroporated with a mixture of sgRNA targeting *PTEN* and PGK-puro plasmids. Cells were cultured at clonal density and selected with puromycin (0.4μg/mL) 48 hr after electroporation for 2 days. Individual hPSC clones were picked, expanded, and genotyped to identify mutant clones. TOPO cloning and plasmid sequencing were used in addition to direct sequencing of the PCR product to confirm the nature of the genetic mutations. To generate *PTEN+ARPC4* double mutant PGP17-iNgn2 hPSCs in a pooled manner, three different clonal *PTEN* mutant PGPC17-iNgn2 hPSC lines (Figure S1) were transduced with lentiviruses containing two different sgRNAs against ARPC4 and selected with puromycin for 5 days (1μg/mL), followed by PCR and Sanger sequencing on genomic DNA. Sequence analysis was performed using the ICE CRISPR Analysis tool (Synthego) to measure gene disruption efficiency.

#### Generation of Ngn2-induced neurons

PGPC17-iNgn2 hPSCs (control, *PTEN* mutant, and *PTEN+ARPC4* double mutant) were cultured in NGD 0.5X medium (45) and treated with 5μg/mL doxycycline (Sigma D9891) on day 1-8. For replating, cells were dissociated with accutase and seeded on day 3 in 0.5X NGD containing doxycycline. 10μM Y27632 was added for the first 24 hours after replating. Depending on the subsequent analyses, replating surfaces include matrigel-coated plates for subsequent immunoblotting analysis, polyethylenimine and matrigel-coated glass coverslips for immunostaining and Phalloidin staining, and polyethylenimine-coated MEA plates for electrical activity measurements. For ARP2/3 inhibitor experiment, differentiating hPSCs were treated with vehicle or CK-666 (100μM) starting on day 1 post doxycycline induction. To sparsely labeling human neurons for morphological analysis, PGPC17-iNgn2 hPSCs were treated with 5μg/mL doxycycline and plated on polyethylenimine-coated glass coverslips on day 3, followed by FUW-G lentivirus transduction on day 7. Neurons were collected on day 14 or day 30 for imaging analysis. To induce neurons from WIBR3 hPSCs (control, *PTEN mutant*, *PTEN+RAPTOR* double mutant and *PTEN+RICTOR* double mutant), dissociated hPSCs were transduced with the pLV-TetO-hNGN2-Neo lentivirus, selected with G418 (200µg/ml) for 5 days and replated on Matrigel-coated plates for doxycycline induction (5µg/ml) in 0.5X NGD media for 3 days. Cells were subsequently replated on the appropriate surface as described above for further assays.

#### Neural precursor cultures

WIBR3 control and *PTEN* homozygous mutant neural precursors (NPs) with complete ablation of PTEN protein were previously generated (9, 10). NPs were maintained in proliferation medium containing 0.5X NGD, 10ng/mL bFGF, and 20ng/mL human insulin on Matrigel-coated plates and passaged using accutase every 5-7 days. For ARP2/3 inhibitor experiment, NPs were treated with vehicle or CK-666 (100μM) for 24 hours. NPs generated from 2 rounds of differentiation or from 2 independent control or *PTEN* mutant WIBR3 hPSC clones were used for a total of 4 biological replicates for each condition. To generate *PTEN+ARPC4* double mutant NPs, *PTEN* mutant NPs derived from two independent hPSC clones were transduced with lentivirus containing Cas9 and sgRNA against scramble or *ARPC4*. The transduced NPs were selected using puromycin (1μg/mL) for 5 days and expanded. Two separate transductions were performed for a total of 4 biological replicates for each genotype condition. The puromycin-selected NPs were harvested for genotype analysis using PCR and Sanger sequencing. Sequence analysis was performed using the ICE CRISPR Analysis tool (Synthego) to measure gene disruption efficiency. Subsequently the same samples were used for immunoblotting analyses to evaluate the protein disruption efficiency.

#### EdU click-it assay

EdU was added to 0.5X NGD medium containing 10ng/mL bFGF and 10ng/mL insulin for NPs for 2 hours, after which cells were fixed and collected. EdU click-it assay was performed on fixed cells per manufacture’s instruction (Thermo Fisher C10458) followed by immuno-staining.

#### Immuno-fluorescent staining, Phalloidin staining, and imaging

For immunostaining, cells were fixed with 4% (w/v) paraformaldehyde (Cedarlane, 15710) in 1X PBS at room temperature for 10 mins. Following membrane permeabilization with PBS containing 0.3% Triton x-100 (Sigma, T8787), cells were blocked with 3% normal donkey serum (Sigma, S30-M). Primary antibodies were against SOX2, KI67, MAP2 and visualized by secondary antibodies conjugated with Alexa 488, 594, 647 (Thermo) followed by counter-staining with DAPI (Thermo, D3571). For Phalloidin staining, cells were fixed with 4% (w/v) paraformaldehyde in PBS and treated with PBS containing 0.3% Triton x-100 (Sigma, T8787) for membrane permeabilization. Cells were blocked with 1% BSA for an hour and stained with Fluorescent phallotoxins (Alexa 488 labelled Phalloidin; Thermo fisher #A12379) for 30 minutes at room temperature according to manufacturer’s protocol. For co-staining with other cell markers after Phalloidin staining, cells were blocked with 3% normal donkey serum (Sigma, S30-M), followed by primary, secondary antibodies and DAPI staining as described above. Fluorescent images of Phalloidin staining and immuno-staining were captured on a Nikon A1R confocal microscope. Quantifications of EdU, KI67, SOX2, MAP2, and DAPI staining were performed using ImageJ. To image sparsely labeled human neurons for morphological analysis, neurons were fixed with 4% paraformaldehyde and imaged with Nikon A1R and Leica SP8 confocal microscope. Sholl analysis and total dendritic length was performed using FIJI (ImageJ).

#### Protein purification and immuno-blotting

Total protein was extracted from cells and tissues using 1X RIPA lysis buffer (Sigma-Aldrich 20-188), with the addition of protease inhibitor cocktail (Sigma-Aldrich 11836170001), phosphatase inhibitor cocktail 2 and 3 (Sigma-Aldrich P5726 and P0044). Total protein from the supernatant was measured using BCA protein assay (Thermo Scientific 23225). The quantified lysate was loaded as 10μg per well, immunoblotted using primary antibodies, and visualized with HRP-conjugated secondary antibodies, using a Radiance Plus Chemiluminescent kit (Azure Biosystems, AC2103) in accordance with the manufacturer’s instructions. Membranes blotted for phospho-proteins were stripped and re-probed with antibodies against total proteins. Membranes blotted for other non-phospho-proteins were stripped and re-probed for GAPDH. Values for phospho-proteins were normalized to total proteins, and other non-phospho-proteins were normalized to GAPDH.

#### Electrophysiology

Doxycycline-induced Ngn2 human neurons were dissociated on day 3 and plated 500,000 cells per well on polyethylenimine-coated 24-well multielectrode arrays and analyzed on the Maestro recording system (Axion biosystems). Recordings were performed daily for 5 minutes until at least day 30. The neural signal processing routine was used to generate timestamps for each field potential recorded at each electrode. Representative raster plots were generated from individual wells to depict all spikes during a 100 second period, using the Neural Metric analysis tool. Network bursts were highlight by peaks and purple rectangles to indicate the synchronous spiking across electrodes in the well. Quantitative data was presented as well-based weighted mean firing rate (overall activity) and area under cross-correlation (synchronicity). Data was analyzed based on individual biological replicates, defined as neurons generated from independent experiments from different hPSC lines, or different sgRNA lentivirus transduction experiment followed by doxycycline induction. Multiple wells were plated for each experimental condition (e.g. genotype, treatment), the recording data were averaged to represent one biological replicate.

### Quantification and statistical analysis

All data values were presented as mean +/-SEM. Student’s t tests were applied to data with two groups. ANOVA analyses were used for comparisons of data with greater than two groups. Post hoc multiple comparison group comparisons were performed with Tukey test. A value of *p*<0.05 was considered significant.

## Supplemental Information

### Inventory of Supplemental Information

**Figure S1.**
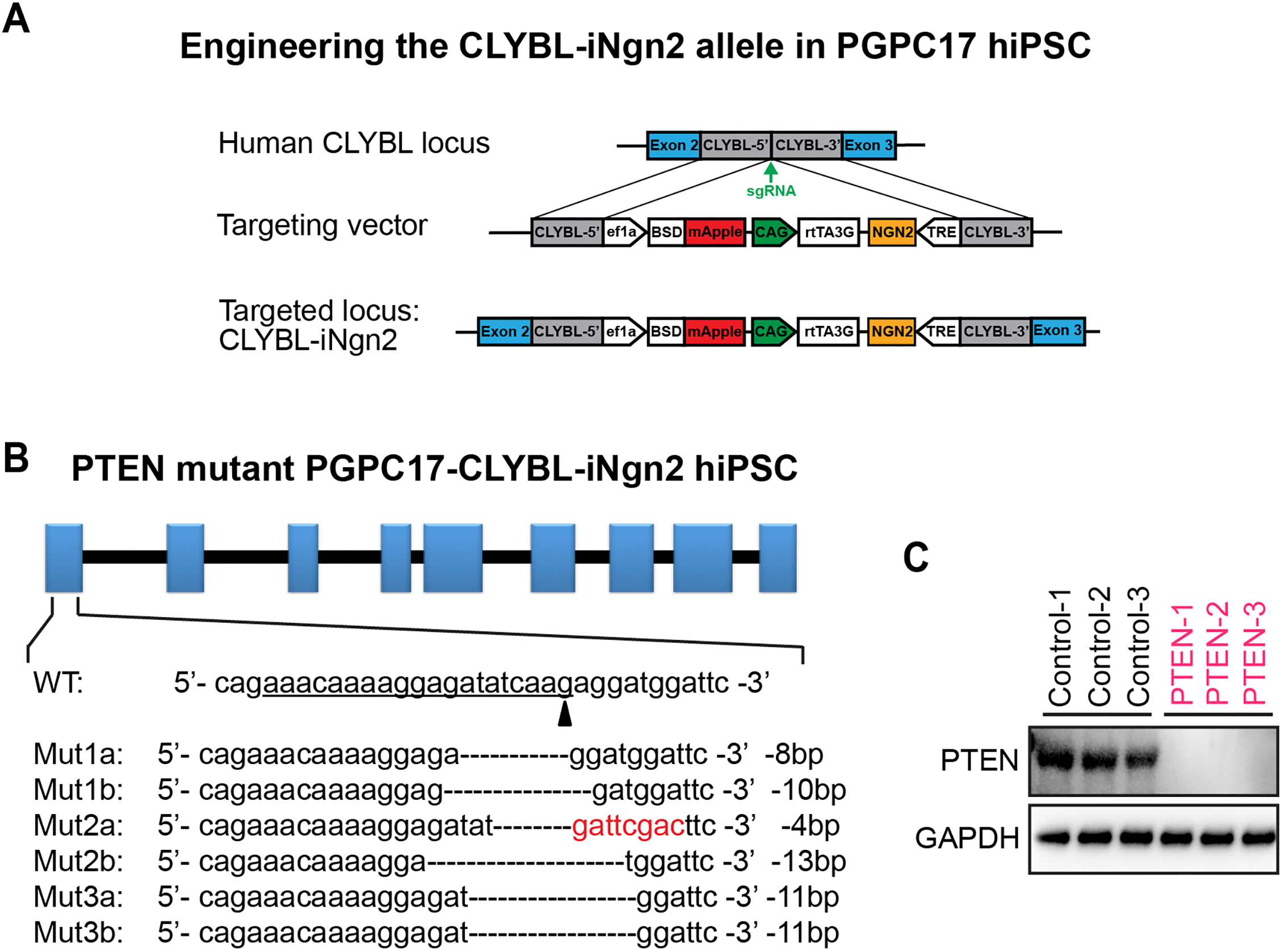
Generating *PTEN* mutant PGPC17 hPSCs. (A) Diagram of the CRISPR-mediated targeting strategy to engineer the iNgn2 containing *CLYBL* locus in PGPC17 hPSC. (B-C) CRISPR-mediated targeting of the human *PTEN* locus, sequences of the mutant PGPC17 hPSC clones (B) and immunoblotting for PTEN protein (C).

**Figure S2.**
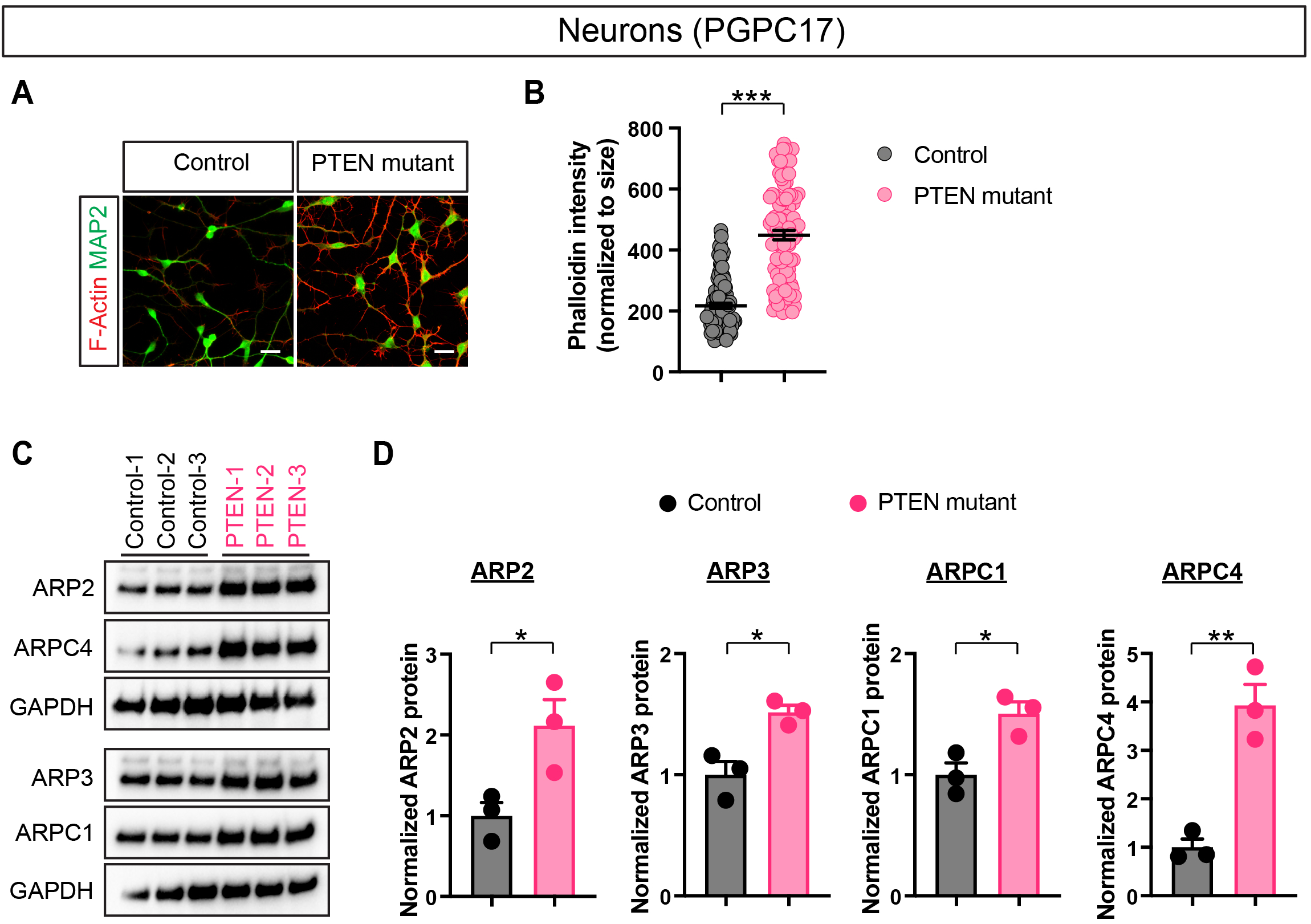
*PTEN* mutant PGPC17 human neurons show increased levels of F-actin and ARP2/3 complex. (A-B) Representative images (A) and quantification (B) of Phalloidin staining show that day 30 *PTEN* mutant PGPC17 human neurons have increased levels of F-actin. Scale bars, 20 μm. Each data point represents a cell (n=100-106 for each group from 3 independent experiments). *p* values are calculated using post hoc multiple comparison after ANOVA. (C-D) Immunoblotting analysis shows that day 14 *PTEN* mutant PGPC17 human neurons have increased levels of key components of the ARP2/3 complex (ARP2, ARP3, ARPC1, and ARPC4). Neurons are induced from PGPC17 hPSCs using Ngn2 induction. Each data point represents an independent experiment (different hPSC clone, n=3 for each group). *p* values are calculated using post hoc multiple comparison after ANOVA. Results are mean +/-SEM. \**p* <0.05, \*\**p* <0.01, \*\*\**p* <0.005.

**Figure S3.**
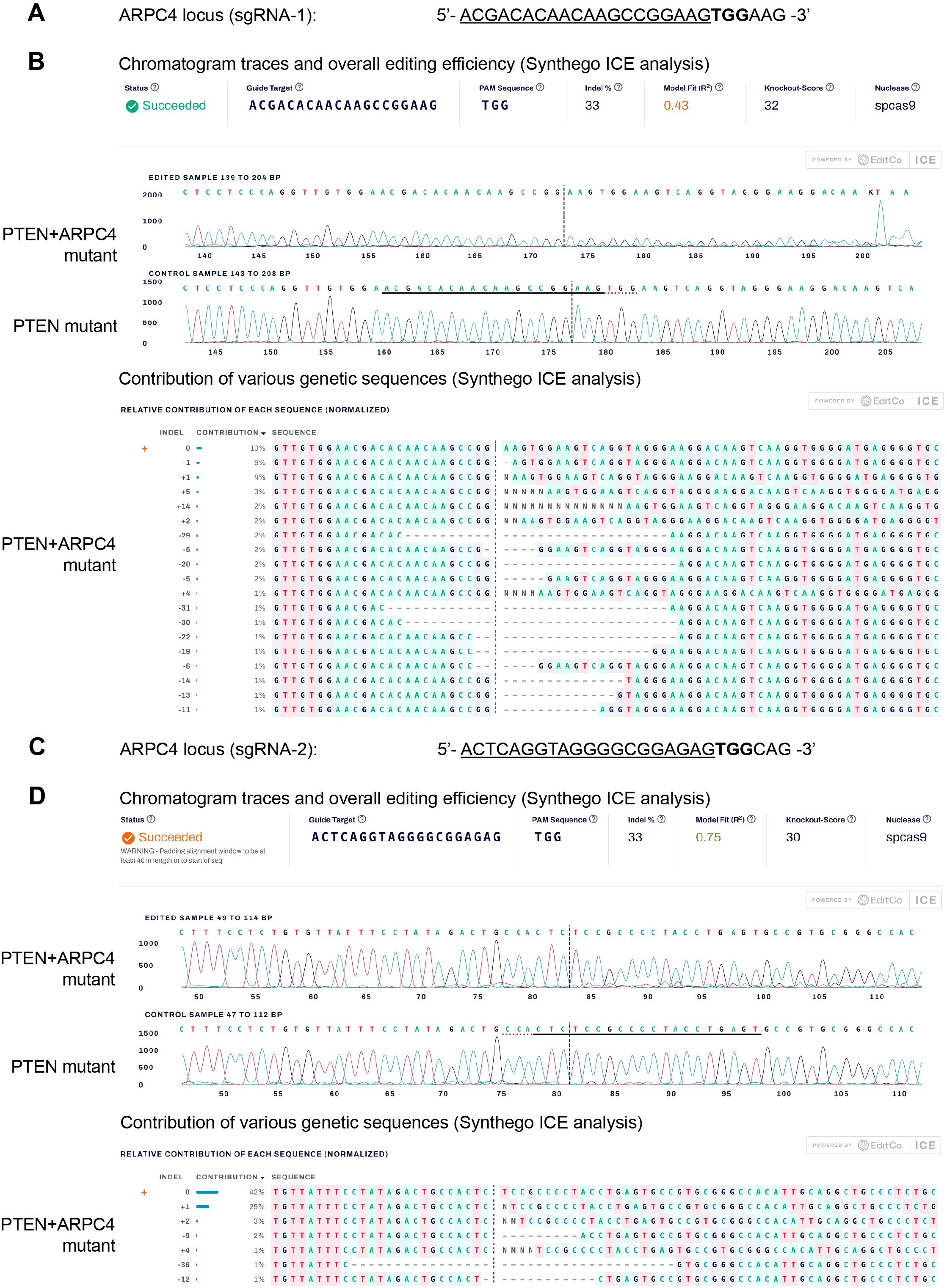
Genetic analysis of the *ARPC4* locus in the *PTEN*+*ARPC4* double mutant WIBR3 human NPs generated in a pooled manner Related to Figure 5. (A) Schematic of the *ARPC4* locus for CRISPR/Cas9 mediated targeting by sgRNA-1. The underlined sequence shows sgRNA-1 sequence and PAM sequence is shown by bold letters. (B) Chromatogram traces, overall editing efficiency, and contribution of individual genetic sequences at the *ARPC4* locus in the *PTEN+ARPC4* double mutant WIBR3 NPs, using Synthego ICE analysis. (C) Schematic of the *ARPC4* locus for CRISPR/Cas9 mediated targeting by sgRNA-2. The underlined sequence shows sgRNA-2 sequence and PAM sequence is shown by bold letters. (D) Chromatogram traces, overall editing efficiency, and contribution of individual genetic sequences at the *ARPC4* locus in the *PTEN+ARPC4* double mutant WIBR3 NPs, using Synthego ICE analysis.

**Figure S4.**
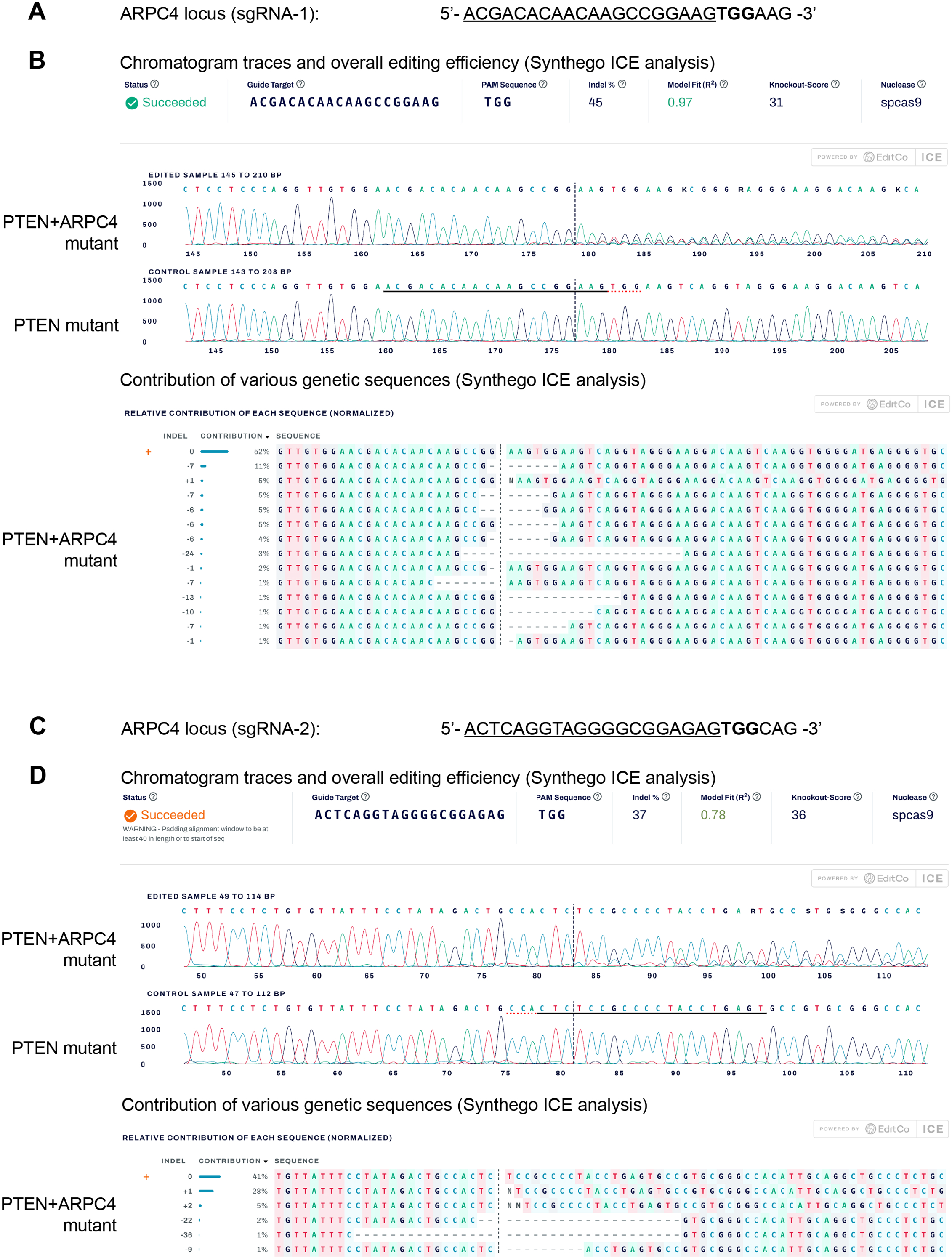
Genetic analysis of the *ARPC4* locus in the *PTEN*+*ARPC4* double mutant PGPC17 human neurons generated in a pooled manner. Related to Figure 6. (A) Schematic of the *ARPC4* locus for CRISPR/Cas9 mediated targeting by sgRNA-1. The underlined sequence shows sgRNA-1 sequence and PAM sequence is shown by bold letters. (B) Chromatogram traces, overall editing efficiency, and contribution of individual genetic sequences at the *ARPC4* locus in the *PTEN+ARPC4* double mutant PGPC17 neurons, using Synthego ICE analysis. (C) Schematic of the *ARPC4* locus for CRISPR/Cas9 mediated targeting by sgRNA-2. The underlined sequence shows sgRNA-2 sequence and PAM sequence is shown by bold letters. (D) Chromatogram traces, overall editing efficiency, and contribution of individual genetic sequences at the *ARPC4* locus in the *PTEN+ARPC4* double mutant PGPC17 neurons, using Synthego ICE analysis.

**Video S1. MEA movie, control and *PTEN* mutant human neurons treated with 100uM CK-666, day 30 post DOX induction.**

**Video S2. MEA movie, control, *PTEN* mutant, and *PTEN*+*ARPC4* double mutant human neurons, day 30 post DOX induction.**

**Table S1.** Antibody information.

**S1a. Antibodies for immuno-staining**
| **Antibody** | **Vendor** | **Catalog #** | **Species** | **Dilution** |
| --- | --- | --- | --- | --- |
| KI67 | Dako | M7240 | Mouse | 1:100 |
| MAP2 | Encor<br>Biotechnology | CPCA-MBP | Chicken | 1:5000 |
| SOX2 | Cell Signaling | 3579 | Rabbit | 1:400 |

**S1b. Antibodies for immuno-blotting**
| **Antibody** | **Vendor** | **Catalog #** | **Species** | **Dilution** |
| --- | --- | --- | --- | --- |
| AKT | Cell Signaling | 9272 | Rabbit | 1:1000 |
| Phospho-AKT (S473) | Cell Signaling | 4058 | Rabbit | 1:1000 |
| ARP2 | ECM<br>Biosciences | AP3861 | Rabbit | 1:1000 |
| ARP3 | ECM<br>Biosciences | AP4581 | Rabbit | 1:1000 |
| ARPC1 | ECM<br>Biosciences | AP4321 | Rabbit | 1:1000 |
| ARPC4 | Everest Biotech | EB08249 | Goat | 1:500 |
| GAPDH | Sigma Aldrich | ABS16 | Rabbit | 1:5000 |
| PTEN | Cell Signaling | 9559 | Rabbit | 1:1000 |
| RAC1 | Thermo<br>Scientific | MA5-32928 | Mouse | 1:1000 |
| S6 | Cell Signaling | 2217S | Rabbit | 1:2000 |
| Phospho-S6<br>(S240/244) | Cell Signaling | 5364S | Rabbit | 1:2000 |
| Rabbit anti goat-HRP | Thermo<br>Scientific | 31402 | Rabbit | 1:5000 |
| Goat anti rabbit-HRP | Thermo<br>Scientific | 31460 | Goat | 1:10000 |
| Goat anti mouse-HRP | Thermo<br>Scientific | 31430 | Goat | 1:10000 |

**Table S2.**
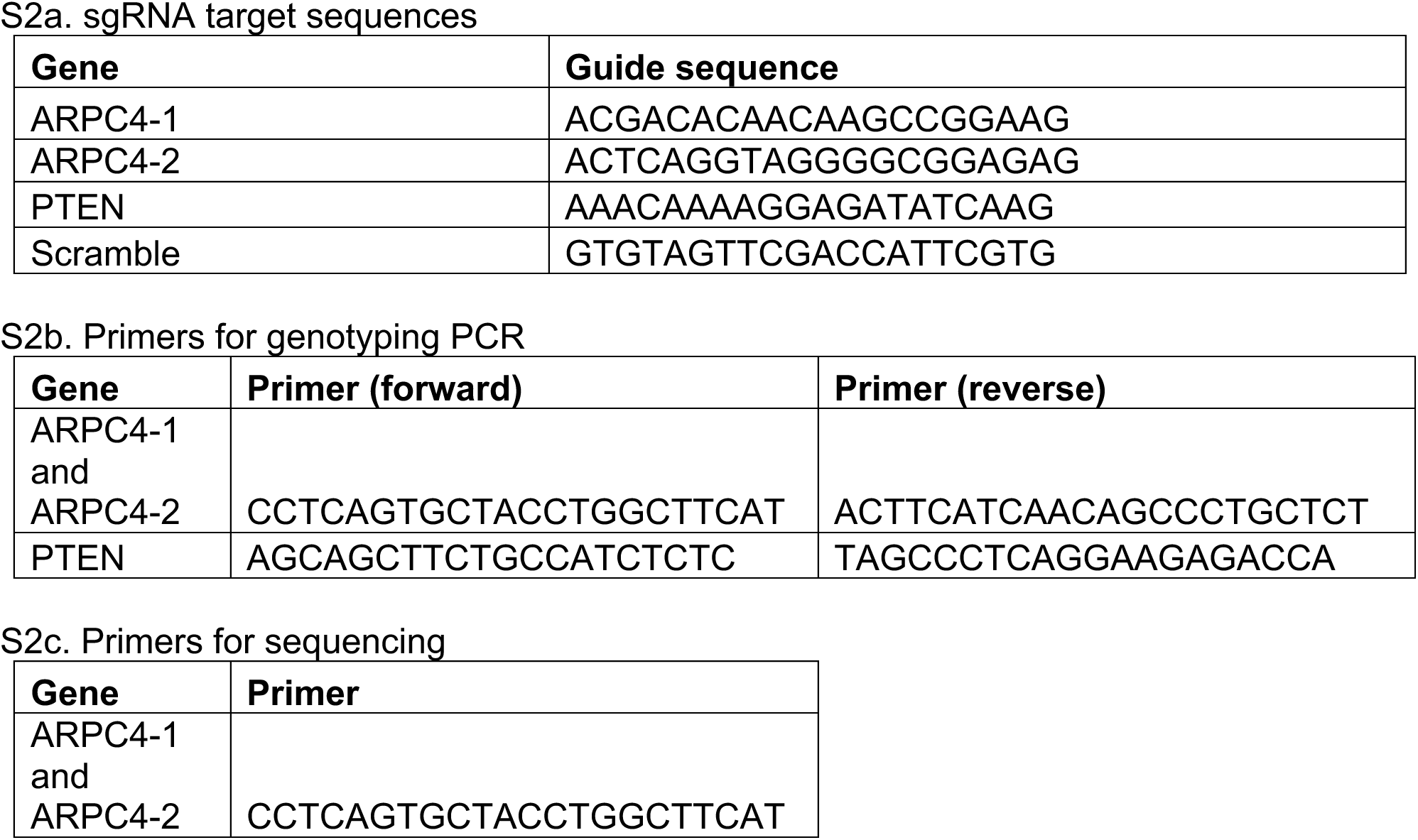
Oligonucleotide information.

